# Acrylic Acid Modified Poly-ethylene Glycol Microparticles for Affinity-Based release of Insulin-Like Growth Factor-1 in Neural Applications

**DOI:** 10.1101/2024.09.25.614803

**Authors:** Pablo Ramos Ferrer, Shelly Sakiyama-Elbert

**Affiliations:** Department of Chemical Engineering, University of Washington, Seattle WA, USA; Department of Bioengineering, University of Washington, Seattle WA, USA

## Abstract

Sustained release of bioactive molecules via affinity-based interactions presents a promising approach for controlled delivery of growth factors. Insulin-like growth factor-1 (IGF-1) has gained increased attention due to its ability to promote axonal growth in the central nervous system. In this work, we aimed to evaluate the effect of IGF-1 delivery from polyethylene-glycol diacrylate (PEG-DA) microparticles using affinity-based sustained release on neurons. We developed PEG-DA-based microparticles with varying levels of acrylic acid (AA) as a comonomer to tune their overall charge. The particles were synthesized via precipitation polymerization under UV light, yielding microparticles (MPs) with a relatively low polydispersity index. IGF-1 was incubated with the PEG-DA particles overnight, and formulations with a higher AA content resulted in higher loading efficiency and slower release rates over 4 weeks, suggesting the presence of binding interactions between the positively charged IGF-1 and negatively charged particles containing AA. The released IGF-1 was tested in dorsal root ganglion (DRG) neurite outgrowth assay and found to retain its biological activity for up to two weeks after encapsulation. Furthermore, the trophic effect of IGF-1 was tested with stem cell-derived V2a interneurons and found to have a synergistic effect when combined with neurotrophin-3 (NT3). To assess the potential of a combinatorial approach, IGF-1-releasing MPs were encapsulated within a hyaluronic acid (HA) hydrogel and showed promise as a dual delivery system. Overall, the PEG-DA MPs developed herein deliver bioactive IGF-1 for a period of weeks and hold potential to enable axonal growth of injured neurons via sustained release.

## Introduction

The locomotor function of the central nervous system is lost after spinal cord injury (SCI), when the axonal pathways extending from the brainstem are severed. Targeting these damaged neuronal circuits and promoting their regeneration and regrowth is a promising therapeutic approach. IGF-1 is a growth factor implicated in cellular function, metabolic homeostasis and most notably for regulating growth and increasing cell survival^1–5^. In the CNS, IGF-1 has been shown to play a role in neuroprotection and has a trophic effect, attenuating motor neuron apoptosis and cell death^6^. Recent studies have shown that the combination of IGF-1 and osteopontin increases corticospinal tract axon growth in a complete SCI model in adult rats^7,8^. The effect of IGF-1 is believed to be mediated mainly via the IGF-1 receptor (IGF-1R), which is highly expressed in neurons, and is upstream of the PI3-kinase/Akt pathway^9^. This receptor also acts as a modulator of signaling endosome trafficking, and studies have shown that IGF-1R activation plays a critical role in growth cone assembly and therefore is important in axonal formation^10^. It also has been reported to mediate enriched environment effects during visual cortical development^1^. The downregulation of IGF-1R in mature neurons and the decreased growth factor levels following SCI are implicated in the limited regenerative ability of adult mammals. IGF-1 has an isoelectric point of 8.2 and a molecular weight of 7.6 kDa^11^. Because of its neuroprotective effect, controlled release of IGF-1 presents a promising target for axon growth if delivered in a sustained manner and a bioactive form.

Biomaterials, including hydrogels and particulate systems, have been extensively explored as drug carriers to the spinal cord ^12–15^. Their high degree of tunability allows for precise control over the material properties, serving as a delivery system for growth factors and other drugs. Loading of molecules into carriers can occur by encapsulation during crosslinking or via incubation and adsorption. In the first case, during delivery system fabrication, growth factors are often exposed to conditions including organic solvents, extended light exposure, sonication, and other stresses.

This exposure may lead to denaturation of the proteins, resulting in decreased biological activity. Prioritizing bioactivity in drug delivery systems will enable more effective therapies and allow for lower loading concentrations, resulting in fewer unwanted side effects and lower cost. Loading of neurotrophic factors onto poly(lactic-coglycolic acid) (PLGA) nanoparticles via short-range electrostatic interactions enabled adsorption and long-term release of the proteins without encapsulation, taking advantage of affinity binding^16,17^. PEG-DA microparticles have been previously synthesized using precipitation photopolymerization ^18^. Both visible and UV light were used to crosslink the polymers, forming small particles with relatively high aggregation but low polydispersity. Furthermore, the zeta potential of the microparticles could be controlled by varying the content of the comonomer, AA. PEG-DA hydrogels have also shown promise as a growth factor delivery platform due to its tunability and properties. Their efficiency can be enhanced by incorporating bioactive peptides and exploiting reversible, affinity binding with growth factors and other proteins, or even covalently tethering proteins to the gels ^19–25^. Previous work from our group has demonstrated affinity binding and sustained release of bioactive NT3 from HA hydrogels ^26^. The charge differences between the carboxylates on the HA chain and the positively charged NT-3 enabled affinity binding of the NT-3 and controlled release of the growth factor over 4 weeks. The hydrogels matched the mechanical properties of native rat spinal cord tissue and allowed adhesion and growth of DRG neurons and mouse embryonic stem cell (ESC)-derived motoneurons and V2a interneurons.

In this work, we explored the potential of modifying PEG-DA based MPs to enable sustained release of IGF-1 by taking advantage of electrostatic interactions between the MPs and cationic proteins. First, we investigated the effect of copolymerization of PEG-DA MPs with AA on IGF-1 loading and delivery. The particles were also embedded within HA scaffolds to further characterize the release profile. Finally, the cytotoxicity and biological activity of IGF-1 were tested using primary neurons and mouse ESC-derived interneurons *in vitro*, as well as in combination with NT-3 as a potential approach to increase axonal growth and neuronal regeneration.

## Materials & Methods

### Polyethylene Glycol-Diacrylate (PEG-DA) microparticle synthesis

PEG-DA MPs were synthesized via precipitation photopolymerization of PEG-DA ^18^ with certain modifications. 3% w/v PEG-DA (MW 3500, VWR), 10 μM of lithium phenyl(2,4,6-trimethylbenzoyl) phosphinate (LAP) (TCI chemicals) as a photoinitiator, AA (Sigma Aldrich) as a comonomer at 0, 40 or 80 mol% relative to PEG-DA and 500 mM of sodium sulfate (Sigma Aldrich) were combined at room temperature. The mixed solution was exposed to UV light at 2.5 mW/cm^2^ and a wavelength of 365 nm for 30 seconds. Following polymerization, the MPs were buffer exchanged to remove the remaining salt in solution via centrifugation at 8000 rpm for 5 minutes.

### Microparticle characterization

10 µL of MP solution was placed on top of a microscope slide and covered with a glass coverslip to reduce motion and drying of the samples. Images were taken at 4x, 10x and 20x magnification in a Nikon Ti2 Microscope using phase contrast imaging. Particle size was determined using ImageJ by measuring the average diameter of each individual MP for at least 10 representative images in 3 distinct replicates.

Polydispersity index (PDI), as a measure of quality with respect to size distribution, was calculated using the following equation:

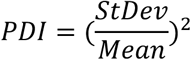

For PDIs under 0.1, a particle solution is assumed to be monodisperse.

### Cytotoxicity of PEG-DA microparticles

V2a interneuron aggregates were prepared as previously described ^27,28^. Briefly, mouse ESCs were differentiated using a 2^-/^4^+^ induction protocol, culturing approximately 1.5 x 10^6^ cells in suspension in agar-coated petri dishes using DFK5 media, composed of DMEM/F12 (Life Technologies) with 5% Knockout Serum Replacement (Life Technologies), 1x Insulin-Transferrin-Selenium (Life Technologies), 50 μM nonessential amino acids (Life Technologies), 5 μM thymidine, 100 μM beta-mercaptoethanol (Sigma) and 15 μM of the following nucleosides: adenosine, cytosine, guanosine, and uridine (Life Technologies). As part of this process, mESCs aggregate into embryoid bodies (EBs), with media changes at two days adding 10 nM RA and 1 µM purmorphamine (EMD Millipore), and at day 4 with DFK5 supplemented with 10 nM RA, 1 μM purmorphamine and 5 μM N-{N-(3,5-difluorophenacetyl-l-alanyl))-(S)-phenylglycine-t-butyl-ester (DAPT; Sigma). At day 6, EBs were dissociated using 0.25% Trypsin-EDTA and quenched with complete DFK5 media. After counting, approximately 3.5 x 10^6^ cells were seeded onto laminin-coated T25 flasks in “selection media”, composed of half DFK5 and half neurobasal (NB) media supplemented with 4 μg/mL puromycin, 10 ng/mL of NT-3, brain derived neurotrophic factor (BDNF) and glial cell line-derived neurotrophic factor (GDNF), 1x GlutaMAX and 1x B27. Two days after selection, motoneurons and V2a interneurons were lifted from the flasks using Accutase ® (Sigma) treatment for 20 minutes and 600,000 – 800,000 cells were transferred into a single well in a Aggrewell 400 (Stemcell Technologies) plate with 1200 microwells per well to get approximately aggregates with 500 – 600 cells per neuroaggregates. The well had been previously treated with anti-adherence rinsing solution to reduce surface tension and reduce cell adhesion. In both cases, the cells were cultured for two days in DFK5:NB media supplemented with 1x GlutaMAX, 1x B27 and 10 ng/mL of NT-3, GDNF and BDNF to drive neuroaggregate formation. At this time, neuroaggregates were lifted and collected for counting before seeding.

PEG-DA MPs with and without AA were synthesized and lyophilized. Following sterilization, 10 mg of particles /1 mL of media were added to each well of a 48 well plate containing 20-30 V2a interneuron aggregates. Samples were run in quadruplicate. DFK5:NB media supported with B-27, GlutaMax and 10 ng/mL NT-3, BDNF and GDNF was used. 24 hours later, a cytotoxicity kit was used to determine the live/dead percentage. Briefly, Calcein AM and ethidium homodimer were added to each well and incubated for 1 hour at 37°C. The wells were gently washed with fresh phosphate buffer saline (PBS) and replaced with fresh media for imaging. Microscope images at 10x were taken of each aggregate in both fluorescence channels. Images were prepared for analysis by splitting the channels and the fluorescent area of live and dead cells was determined in ImageJ using associated functions. First, background signal was removed from images. Using the threshold function, the area in pixel intensity was measured for both excitation channels. The percentage of live cells was calculated by: Area of Calcein-AM stain/ (Area of Calcein-AM + Area of Ethidium homodimer) x 100.

### IGF-1 loading

PEG-DA MPs were prepared as mentioned. Stock solutions of human recombinant insulin (Sigma-Aldrich) and IGF-1 (STEMCELL Technologies) were made in PBS supplemented with 0.1% BSA (Sigma Aldrich) at 20 mg/mL and 1 mg/mL, respectively. Briefly, 10 mg of microparticles per sample were prepared and resuspended in 200 µL of PBS with 100 µg insulin or 10 µg IGF-1. The solutions were allowed to equilibrate for 24 hours at 4°C. For an indirect measurement, 100 µl of loading solution were removed to measure the amount of free protein. The theoretical loading was calculated using the free protein (non-loaded) and the original known mass of protein added to the microparticles. For a direct measurement, the MPs were degraded over two hours with 200 µl dimethyl sulfoxide (DMSO) (Sigma Aldrich) and 600 µl of 0.2 M NaOH (Sigma Aldrich) solution added to each sample. The amount of protein in the microparticles was measured using a Pierce BCA or Micro BCA assay kit (ThermoFisher).

### Long term release of IGF-1 from PEG-DA microparticles

To investigate the drug release characteristics of IGF-1 from PEG-DA MPs, a four-week long release study was conducted. PEG-DA MPs were synthesized as previously mentioned, and 60 mg each of 3 different formulations were prepared: PEG-DA, and PEG-DA with 40 or 80 mol% AA relative to PEG-DA. 15 mg of MPs per sample (all conditions tested in quadruplicate) were incubated in 200 µl PBS with 15 ug IGF-1 for 24 hours at 4°C. Once the growth factor was loaded, microparticle solutions were centrifuged to remove the remaining IGF-1 that had not fully bound to the particles. The samples were then resuspended in 500 µl PBS as a release buffer and 200 µL samples were collected at the following time points: 1, 3, 5, 7, 10, 14, 21 and 28 days. The release buffer was replenished with 200 µL fresh PBS to keep the total volume constant. Following the collection of the last sample, MPs were degraded in 200 µL DMSO and 600 µL of 0.2M NaOH for 6 hours to extract the remaining growth factor. Sample concentrations were measured using a MicroBCA assay kit (ThermoFisher) comparing readings at 562 nm to that of a standard curve.

Alternatively, to understand the mechanism of a potential dual-release system, PEG-DA MPs were loaded with IGF-1 and encapsulated in hyaluronic acid methylfuran (HAmF hydrogels^26,29,30^. PEG-DA microparticles were prepared, incubated for 24 hours with 20 µg IGF-1 to load the growth factor and centrifuged to obtain a pellet. 60 mg each of 3 different formulations were prepared: PEG-DA, PEG-DA with 40 or 80 mol% AA relative to PEG-diacrylate. HAmF hydrogels were prepared by mixing HAmF PEG-dimaleimide at a 3:1 maleimide to methylfuran ratio. The microparticles were added to the HAmF solution before adding the crosslinker. The total volume of each hydrogel was 200 µL. Four separate conditions were tested, one with each MP formulation and one with free IGF-1 added to the gels, all in quadruplicate. The hydrogels were allowed to crosslink and form for 3 hours. Next, 500 µL of PBS were added to each tube as the release buffer and 200 µL samples were collected from this buffer at the following time points: 24 hours, 4 days, 7 days, 10 days, 14 days, 21 days and 28 days. After removing each sample, the buffer was replenished with 200 µL of fresh PBS. At the end of the experiment, the hydrogels were degraded with 2500 U/mL of hyaluronidase, 10 mg/mL heparin and 137 mM NaCl to determine the leftover amount of growth factor. A Micro-BCA assay kit was used to measure the growth factor concentration in each sample by comparing each reading at 562 nm to that of an IGF-1 standard curve of known concentrations. Using these results, a release curve over time was plotted.

### Biological activity of IGF-1 *in vitro*

The bioactivity of IGF-1 was assayed using a neurite outgrowth assay. V2a interneuron aggregates were prepared as described above. At least 15 aggregates were seeded into a well of a 24 well plate, in quadruplicate, using NB:DFK5 media supplemented with 1% GlutaMAX, 1% B27 and the following conditions: 10 ng/mL of NT-3, GDNF and BDNF as the growth factor positive control, or 0, 25, 50, 100 ng/mL IGF-1. For the combination study, 20 ng/mL NT-3 and 50 ng/mL IGF-1 as well as 10 ng/mL NT-3 and 25 ng/mL IGF-1 were tested together to investigate synergistic effects of both growth factors. 2 days after seeding, the aggregates were imaged using fluorescence microscopy using their expression of a TdTomato reporter. The average radius of axonal extension was measured and analyzed on ImageJ software.

### Released IGF-1 Bioactivity on DRGs

Briefly, microparticles were formed with 40% mol AA relative to PEG-DA in a concentration of 3% in neurobasal (NB) media. 250 ng/mL IGF-1 was loaded onto the MPs over 24 hours of incubation at 4°C. Samples of 300 µL were prepared in triplicate and at the selected timepoints, namely 24 hours, 7 days and 14 days following encapsulation of the growth factor, the MPs were centrifuged, and the supernatant collected. For the released IGF-1 bioactivity study, DRGs were harvested from day 7 White Broiler chicken embryos. Approximately 5 DRGs were pooled and placed in a well of a 12 well plate, with at least 4 wells in each condition. Fresh media was prepared using NB supplemented with 1% GlutaMAX, 1% penicillin-streptomycin, 2% B-27 supplement and 0.1 % BSA. For the negative control, no growth factors were added, for the GF control, 10 ng/mL NT-3, GDNF and BDNF were added, and 50 ng/mL IGF-1 was used as the positive control. For the released samples, the collected media with IGF-1 was used and adjusted to the total volume of the NB solution. 48 hours post-seeding, images were taken using phase microscopy. Average neurite extension from the DRGs was measured using ImageJ and normalized to the GF control between experiments.

### Western Blot

V2a interneurons were exposed to 20 ng/mL NT-3, 50 ng/mL IGF-1 or both for one hour. Cell lysates were obtained by trypsinizing and collecting the cells in ice cold RIPA buffer (1 mL per 1x10^7^ cells), centrifuging and collecting the supernatant. The protein concentration was quantified using a bicinchoninic acid (BCA) assay (Pierce) and a dilution of protein concentration standards. Based on the results of the BCA, 20 µg of protein lysate were loaded into each well of a 4% Mini-PROTEAN TGX Stain-Free protein gel (BioRad), and the proteins were separated using gel electrophoresis at 200 mV. The proteins were then transferred to Immun-Blot PVDF membranes (BioRad). The membranes were blocked in EveryBlot Blocking Buffer (BioRad) for 1 h followed by an overnight incubation with primary antibody. Primary antibodies for Erk1/2 and p-ERK1/2 (Cell Signaling Technology), were used at a 1:1000 dilution. The following day, the membranes were washed five times with TBS containing 0.1% Triton-X (TBS-T). The membranes were then incubated with secondary antibody at a 1:10000 dilution (Cell Signaling) for 1 h followed by an additional five washes with TBS-T. The blots were then developed using SuperSignal West Pico Plus chemiluminescence substrate (ThermoFisher) for 5 minutes before imaging on a GelDoc imaging system (BioRad). Molecular weight was determined by comparison to Precision Plus Protein Standards (BioRad). The blots for ERK1/2 and p-ERK1/2 were developed on the same blot by stripping the antibodies with 0.2M glycine in DI water at pH 2.2 with 0.1% Tween-20 for 20 minutes before reapplying the primary antibody.

## Results

### Formation and characterization of PEG-DA microparticles

PEG-DA MPs with or without AA were photopolymerized in the presence of LAP. 500 mM sodium sulfate was added before UV exposure to aid in phase separation, previously found to be the optimal salt concentration^18^. We used Irgacure 2959 as a photoinitiator in previous experiments but found LAP to be more efficient due to its better water-solubility. The MPs were formed rapidly, with less than 30 seconds of exposure to a 365 nm wavelength UV lamp.

There were not visible differences in shape or size for the MPs with changes in AA concentration, even though the conditions with AA tended to aggregate more than those without **(Figure 1a)**. However, this aggregation was easy to disrupt by gently pipetting up and down the MPs in solution. Regardless of the formulation, all MPs ranged in diameter between 2 and 3 µm **(Figure 1b and Table 1).** We did not expect the differences in formulation to significantly alter the size and morphology of the MPs based on our preliminary experiments and previous work^18^.

**Figure 1.**
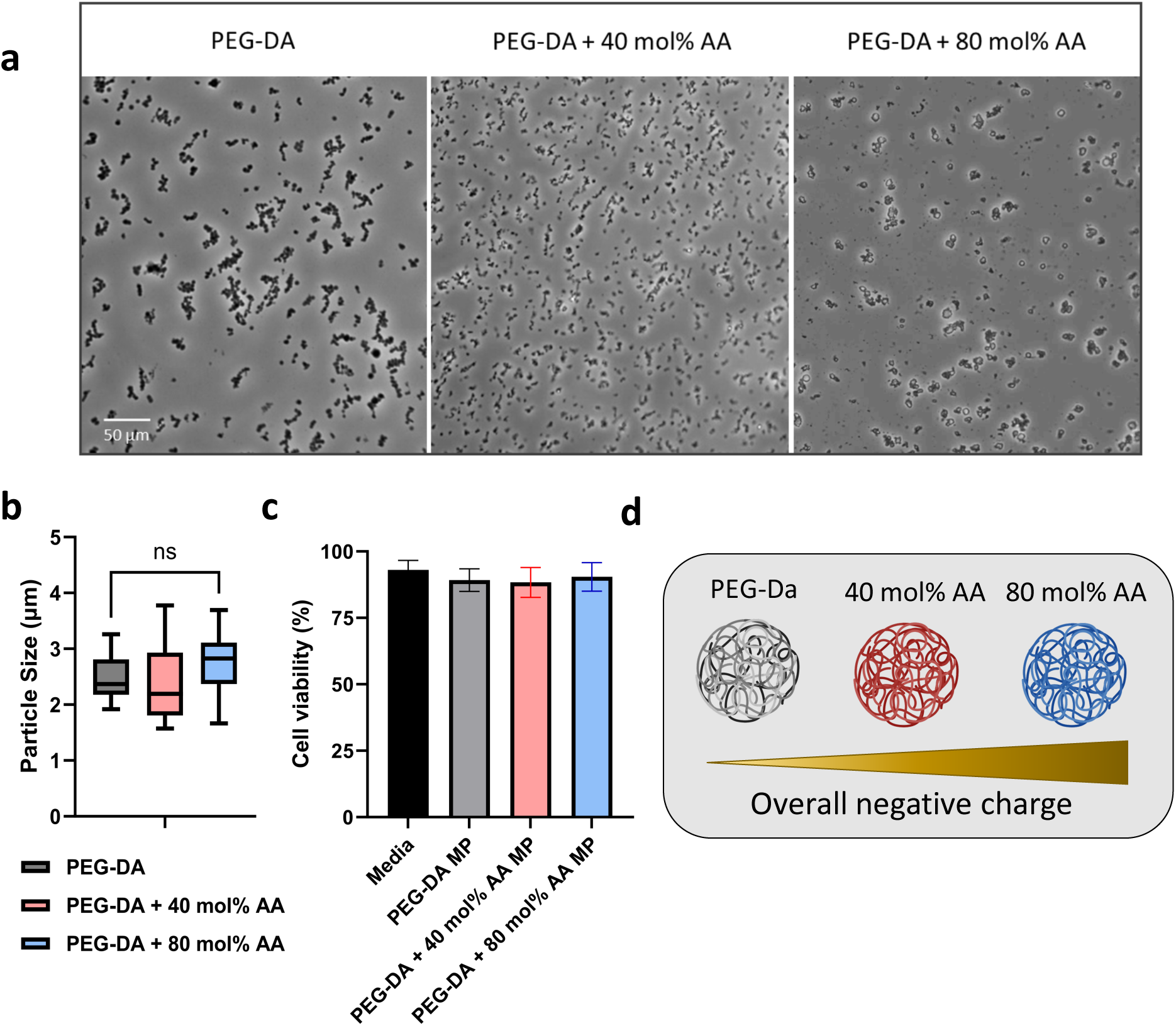
a) Representative images of PEG-Diacrylate microparticles with 0%, 40% or 80% mol acrylic acid relative to PEG-DA in the presence of LAP as a photoinitiator (scale bar = 50 µm). b) Average microparticle size for PEG-Diacrylate with 0%, 40% or 80% mol acrylic acid relative to PEG-DA in the presence of LAP as a photoinitiator (* = p< 0.05, ** = p< 0.01, *** = p< 0.001). c) Cell viability of V2a interneuron aggregates treated with PEG-DA microparticles reported as a percentage of total cells present in each condition. d) Schematic of the PEG-DA microparticles formulations used in the study based on acrylic acid content.

**Table 1.** Average microparticle size and polydispersity index for PEG-diacrylate formulations.

| | Size ($\mu\text{m}$) | PDI |
| --- | --- | --- |
| PEG-DA | 2.327 | 0.071 |
| PEG-DA + 40 mol% AA | 2.292 | 0.043 |
| PEG-DA + 80 mol% AA | 2.795 | 0.022 |

The polydispersity index ranged from 0.071 for PEG-DA to a low of 0.022 in the case of 80 mol% AA relative to PEG-DA in the presence of LAP. All measured values fell within the monodispersed range, with PDI under 0.1. This low degree of polydispersity suggests that the precipitation polymerization reaction produces a highly homogenous mix of particles.

To ensure that our PEG-DA based MPs were biocompatible with cells and specifically neurons, we tested their cytotoxicity using a live/dead assay. In the presence of 10 mg MPs/ 1 mL media, there was no statistically significant difference between the control and the microparticles, independent of the AA content and overall MP formulation **(Figure 1c).** All cell viability percentages in V2a interneuron aggregates exposed to the MPs ranged from 87% to 93%, numbers that confirm the *in vitro* biocompatibility of the PEG-DA MPs and suggest that they could likely be a viable candidate to be implanted *in vivo* without a significant cytotoxic response.

### Loading of insulin via incubation

Following physical characterization, we tested the potential of these PEG-DA based MPs to load proteins. We used insulin as a control protein based on its similar size but different isoelectric point to IGF-1. At pH 7.4, insulin is negatively charged, while IGF-1 is electropositive. We tested all formulations by fabricating, washing and drying the particles, then loading insulin or IGF-1 by incubating the protein with MPs over 24 hours. We then measured the % loaded indirectly, by measuring the remaining unloaded insulin or IGF-1 in solution after spinning down and removing the supernatant, and directly, by degrading the MPs in DMSO and NaOH before reading out the protein concentration. In the case of insulin, we found no clear trend or correlation between the different conditions, with loading concentrations between 37% and 51% for all groups **(Figure 2a).**

**Figure 2.**
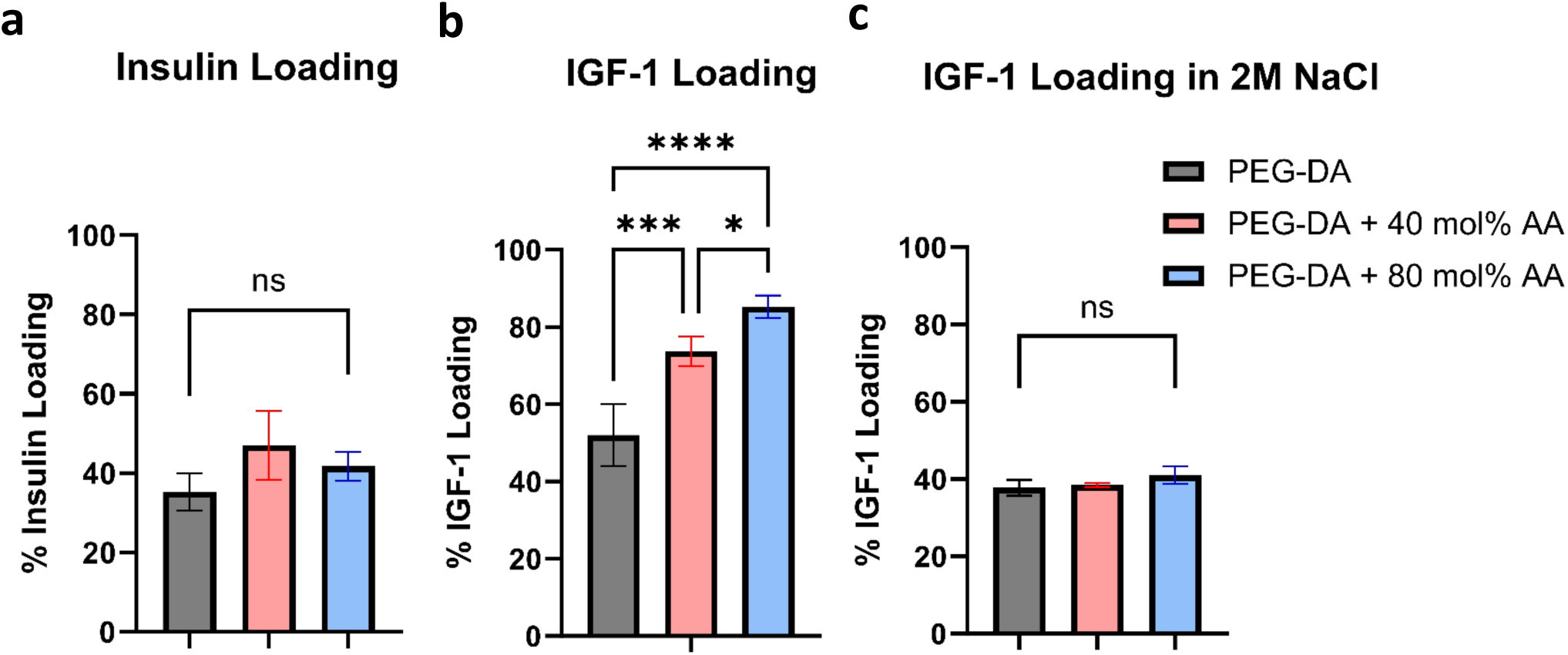
Percentage loading for PEG-diacrylate with 0%, 40% or 80% mol acrylic acid relative to PEG-DA in the presence of LAP as a photoinitiator of a) 100 µg insulin, b) 10 µg IGF-1, and c) 10 µg IGF-1 in a solution containing 2M NaCl per condition. Error bars denote standard error of the mean (* = p< 0.05, *** = p< 0.001, **** = p< 0.0001).

However, when we tested the same MPs with IGF-1 we saw differences in loading in between our three MP formulations. As the amount of AA increased, we observed significantly higher IGF-1 loading, going from ∼50% to over 80% for the group with the high AA concentration **(Figure 2b).** One of the main reasons why this trend was present is the difference in protein properties. Insulin, with an isoelectric point of 5.4 and 5.7 kDa size, and IGF-1 with pI= 8.5 and a molecular weight of 7.6 kDa, present different loading behavior. We attribute this differential loading to the different protein charge in the PBS solution. IGF-1, is positively charged at physiological pH, while the carboxylate groups present in the PEG-DA MPs yield an overall negative charge over the particles. This behavior was not present in the case of insulin, suggesting that its low isoelectric point and negative charge are not sufficient to drive protein encapsulation.

To further characterize this behavior, we loaded IGF-1 in the presence of 2M NaCl to shield the microparticle charges. We found that in this case, the overall amount encapsulated was reduced in all conditions and there were no differences in loading between the MP formulations **(Figure 2c).** More specifically, in the case of the 80 mol% AA condition, the fraction of IGF-1 loaded decreased from 0.85 to 0.4, confirming that a high salt concentration disrupts the electrostatic interactions present. Using a high salt concentration effectively neutralized the charge differences between each of the formulations and equalized the final loading percentage regardless of AA content. Therefore, this particulate system provides a tunable loading platform for positively charged or cationic proteins by exploiting affinity binding.

### Release of IGF-1 from PEG-DA microparticles

Once we confirmed the MPs’ ability to load growth factors, we investigated the release profile of IGF-1 over a period of 4 weeks. As expected, based on the loading experiments, we observed slower protein release as the AA concentration increased **(Figure 3a).** Less than 20% of growth factor released after 24 hours in all conditions, and we observed no significant burst release with the MPs displaying sustained release. In the case of PEG-DA without AA, almost 60% of the loaded IGF-1 was released in the first 7 days. However, when 40 or 80% mol AA (relative to PEG-DA) was used, that number dropped to 50 or 40% released, respectively. During the remainder of the release study, IGF-1 was slowly delivered reaching 80% and 60% of the total amount loaded in both 40 and 80% mol AA formulations. In this case, similar to what we observed during encapsulation, we found a direct correlation between binding and AA content, suggesting the presence of stronger interactions when the charge difference between MPs and IGF-1 is greater.

**Figure 3.**
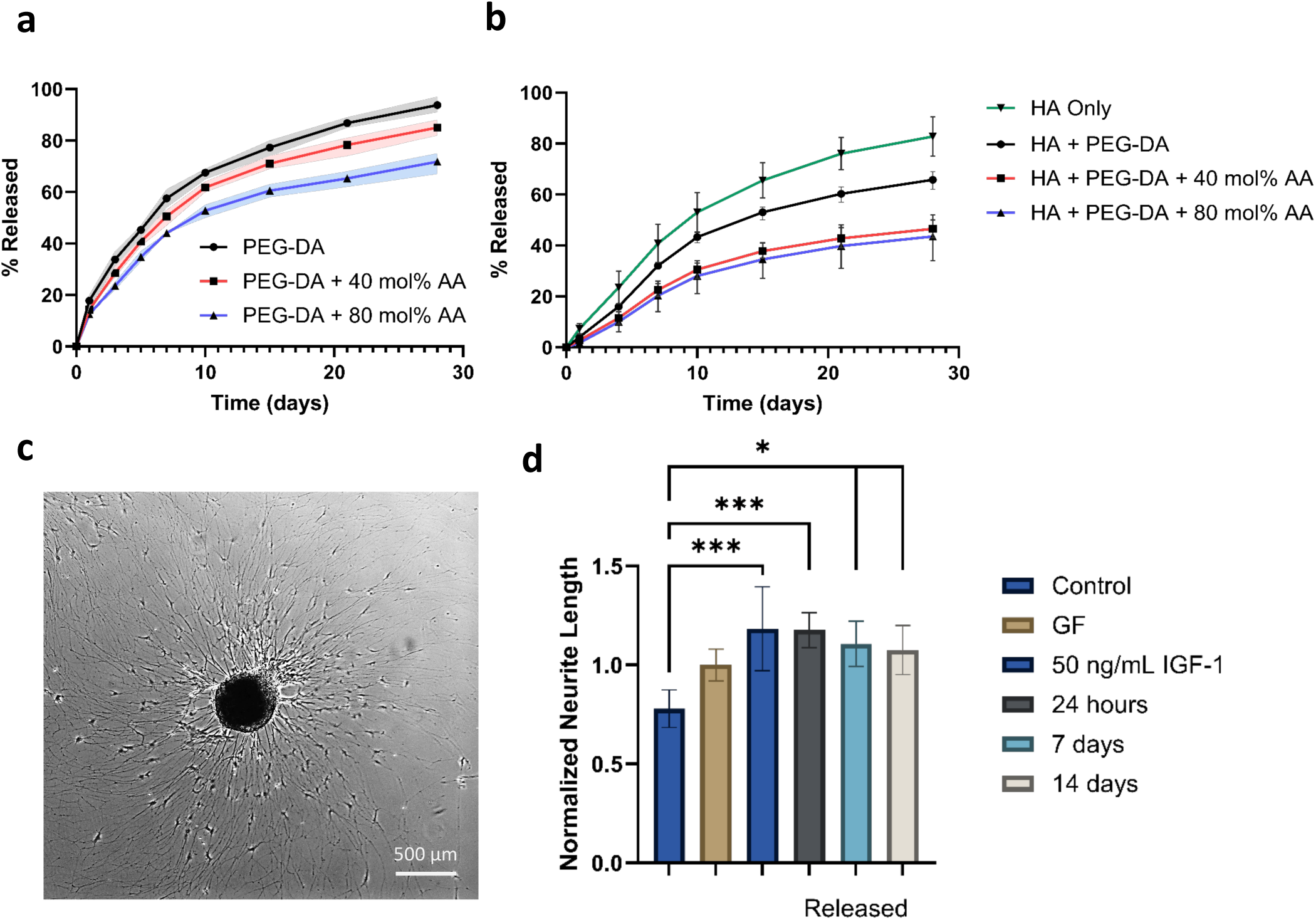
a) Release curves of IGF-1 from PEG-Diacrylate microparticles with 0%, 40% or 80% mol acrylic acid relative to PEG-DA. Each condition contained 15 mg of microparticles and was previously loaded with 15 µg of IGF-1 that was released into PBS buffer (n=4). b) Release curves of IGF-1 released from PEG-Diacrylate microparticles with 0%, 40% or 80% mol acrylic acid relative to PEG-DA embedded in a HA hydrogel. Each condition contained 15 mg of microparticles and was previously loaded with 15 µg of IGF-1 that was released into PBS buffer (n=4). c) Representative image of DRG in the presence of 50 ng/mL fresh IGF-1. d) Average neurite outgrowth from DRGs in the presence of 10 ng/mL NT-3, GDNF, BDNF (growth factor (GF) positive control), 50 ng/mL IGF-1, released IGF-1 from PEG-DA MPs at 24 hours, 7 days and 14 days normalized to the GF positive control (* = p< 0.05, *** = p<0.001, **** = p<0.0001, n=3 for at least 10 neuroaggregates per group).

We then aimed to combine the PEG-DA MPs with a HA hydrogel system to assess the feasibility of a dual-delivery platform. MPs under the same conditions as in the solution study were prepared and encapsulated within the HA hydrogels via incubation during crosslinking at 37°C. Hydrogels were placed in excess PBS buffer, and the concentration of released IGF-1 was determined at each time point. As expected, encapsulating the MPs within the scaffold drove down the release rate in all three formulations **(Figure 3b)**. In the case of only PEG-DA, the total amount released at the end of the 4 weeks decreased to just over 60%, and for both AA conditions the final amount delivered was ∼40% of the loaded IGF-1. In the case of just HA, the growth factor had a faster release and 80% of its initial concentration was delivered by the end of the study. Physically, the entrapment of the particles within the hydrogel pores is one of the drivers of the decreased release rate, and the addition of the electronegative microparticles drives electrostatic binding in the negatively charged scaffold further slows down the delivery of growth factor.

To confirm the biological activity of released IGF-1, we used a DRG neuron bioactivity assay. Whole DRGs from chick embryos were harvested and seeded in laminin-coated wells and supplemented with NB media containing IGF-1 delivered from the PEG-DA MPs. DRGs were exposed to released IGF-1 at 24h, 7 days or 14 days and the neurite length was analyzed. Fresh IGF-1 at a concentration of 50 ng/mL was used as a positive control for comparison with the released IGF-1 and 0 ng/mL IGF-1 was used as the negative control. All measurements were normalized to a growth factor cocktail control, containing 10 ng/mL each of NT-3, BDNF and GDNF. At the 24-hour time point, the average neurite length was almost the same as that of fresh IGF-1. Based on the initial loading of 250 ng/mL and ∼20% of it being released based on the release curves, the IGF-1 appears to retain its bioactivity on the first day of release, and most of it by days 7 and 14 **(Figure 3c, 3d)**. The growth elicited by the release samples from day 14 showed statistically significant difference with the control and no difference with the fresh, 24 hour and 7 day samples. The amount of IGF-1 loaded was adjusted based on release curves for the release samples to have the same concentration at all three time points. Since a therapy of this type would be the most beneficial during the subacute phase of SCI, sustained release over two weeks would be a promising approach to enhance and promote axonal regeneration.

### Biological activity of IGF-1 on V2A interneurons

We then aimed to study the effect of IGF-1 on mouse ESC-derived V2a interneuron aggregates. DRGs are peripheral sensory neurons and do not accurately represent the neuron population present on the spinal cord, so utilizing these types of neurons yields a more relevant result for spinal cord neurons. Several conditions were prepared from 0 to 100 ng/mL IGF-1 to test the cells response to the growth factor, aiming to establish a correlation between IGF-1 concentration and neurite extension **(Figure 4a).** The neurite extension was higher at 10 and 25 ng/mL IGF-1 but not significantly different from the control, and it peaked at 50 ng/mL. The growth at 100 ng/mL was lower than 50 ng/mL, suggesting that the cells response to IGF-1 is biphasic with respect to concentration. This is in agreement with other studies reporting a biphasic response to neurotrophic factors, where a specific dosage yields an optimal response and a higher concentration results in a reduced response, potentially through the saturation of the growth factor cell surface receptors^31,32^.

**Figure 4.**
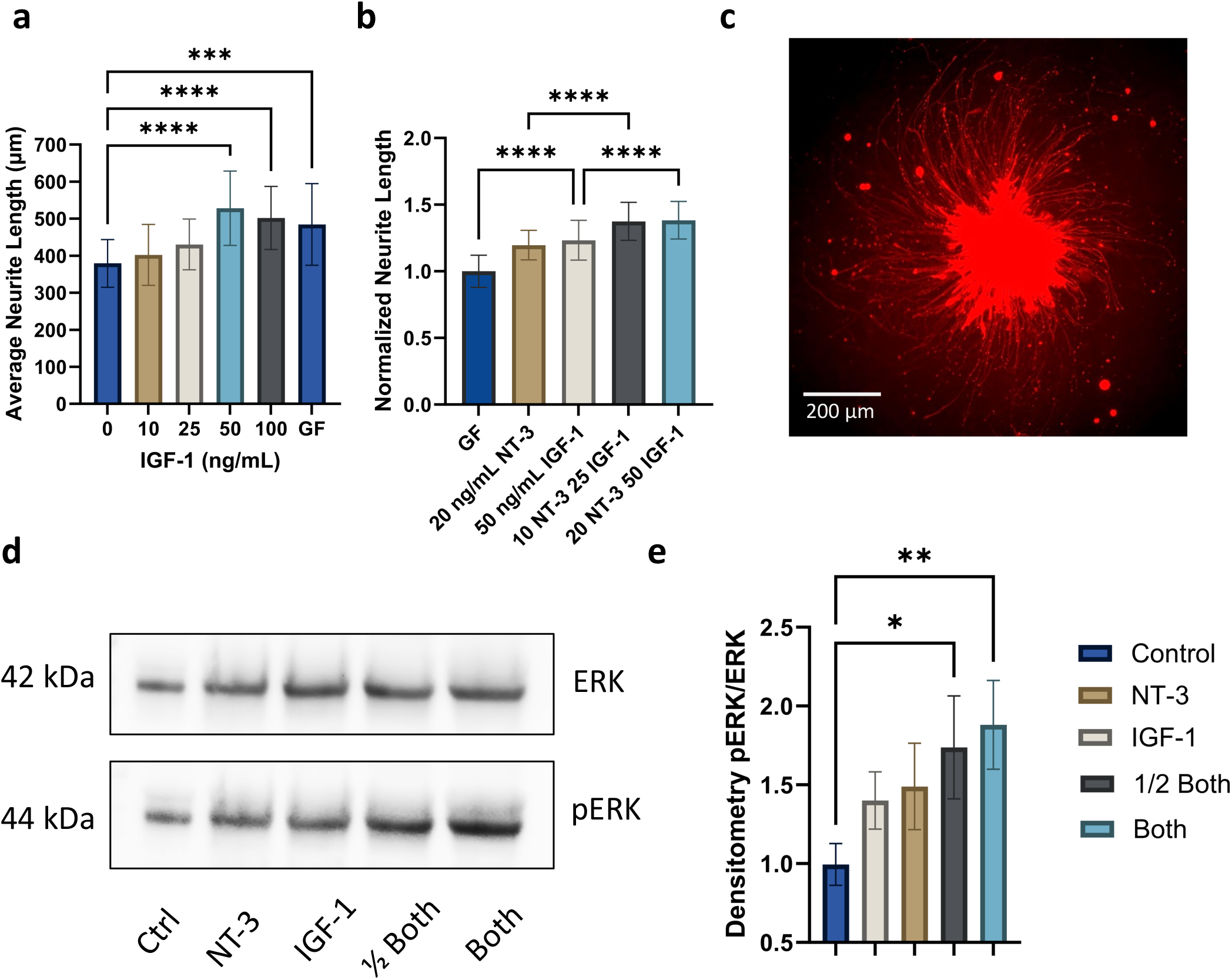
a) Average neurite outgrowth from V2a interneuron neuroaggregates in the presence of 0, 10, 25, 50 or 100 ng/mL IGF-1. (*** = p<0.001, **** = p<0.0001, n=3 for at least 30 neuroaggregates per group). b) Average neurite outgrowth from V2a interneuron neuroaggregates in the presence of 20 ng/mL NT-3, 50 ng/mL IGF-1 or both, normalized to the GF positive control (*** = p<0.001, **** = p<0.0001, n=3 for at least 30 neuroaggregates per group). c) Representative image of tdTomato positive V2a interneuron aggregate in the presence of 50 ng/mL IGF-1. d) Phosphorylation of ERK1/2 in V2a interneuron cell lysates previously exposed to growth factors measured via Western Blot. e) Densitometry of Western Blot data calculated as pERK/ERK and normalized to the control. N=3 (* = p< 0.05, ** = p< 0.01).

Next, we explored whether the combination of IGF-1 with NT-3 would be capable of improving the axonal growth that each of the growth factors could achieve individually. For this, we used V2a interneuron aggregates and added the growth factors concentrations that yielded the highest growth in each of the respective studies, 20 ng/mL for NT-3 and 50 ng/mL in the case of IGF-1 **(Figure 4c)**. We found that the combination of both molecules resulted in a significantly higher neurite length than each of them separately **(Figure 4b).** We also observed a similar enhancing effect of the combination even when both concentrations were cut in half, using 10 ng/mL NT-3 and 25 ng/mL for IGF-1. This suggests the presence of a synergistic effect between the growth factors when acting in combination with each other. Even though IGF-1 signals through the IGF-1 receptor, a tyrosine kinase, and NT-3 mainly through TrkC, a tropomyosin receptor, both growth factors share the downstream signaling pathways phosphoinositide-3 kinase (PI3K)-AKT and RAS-mitogen activated protein kinase (MAPK)^33–35^. We hypothesized that sharing these common pathways plays a role in the synergistic effect observed in the combined treatment in neurons. The negligible difference between the 20 - 50 ng/mL and 10 - 25 ng/mL can be attributed to saturation of these common signaling pathways.

To further investigate this behavior, we measured phosphorylation levels of ERK 1/2, extracellular-signal regulated kinases 1 and 2^36^. Based on previous data from our group, we expected higher phosphorylation levels within 60 to 90 minutes after growth factor exposure^37^. For this reason, we cultured V2a interneurons and exposed them to NT-3, IGF-1 or both for 60 minutes and then collected the cell lysate to measure protein expression via Western Blot **(Figure 4d)**. Densitometry calculations based on band intensity after protein loading normalization showed significantly higher ERK phosphorylation for both conditions in which NT-3 and IGF-1 were combined **(Figure 4e)**. This data suggests that both growth factors share a downstream signaling pathway and the activation of this pathway, likely via TrkC and IGF1-R activation. Phosphorylation of ERK was increased by IGF-1 and NT-3 stimulation, highlighting the potential of the combination of these growth factors as an axonal growth approach, which had been reported previously^38–41^.

## Discussion

Regrowth of lost axonal circuitry has long been a promising therapeutic approach to promote functional recovery following SCI. Insulin-like growth factor-1 has been shown to have a trophic effect, increasing axonal growth both *in vitro* and *in vivo* and even improving locomotor function scores in preclinical models of SCI in rodents ^8,42^. Even though many potential therapeutic agents have been extensively explored, there is still a need to optimize the drug delivery systems employed for SCI. Delivering a sustained concentration of bioactive drug over an extended period of time remains a challenge in the field. Microparticles provide a highly tunable platform to have temporal and spatial control over the release rate of therapeutic agents. Parameters such as size, charge, material choice, degradability and biocompatibility are all critical when designing and developing drug release systems. Here, we can exploit non-specific binding mechanisms to control loading and release, namely electrostatic interactions.

The precipitation polymerization of PEG-DA reported herein produced microparticles of similar size and polydispersity regardless of formulation, with changes in AA content not significantly affecting the average diameter size. Based on previous data, the main size determinant parameters that could be exploited are the molecular weight of the initial PEG-DA, and the concentration of Na_2_SO_4_^18^. The underlying model that describes the mechanism of nucleation and production of monodisperse particles was first reported 70 years ago^43^. This model set up the theoretical basis for other microparticle production methods, including the precipitation polymerization used here. According to La Mer, particle formation occurs in two steps: nucleation and growth. In the initial nucleation step, phase separation occurs when small clusters of atoms or molecules assemble to form a nucleus. Supersaturation of the polymer is necessary for nucleation to occur, and we controlled this by adding relatively high salt concentrations. Growth of newly formed particles proceeds in the absence of nucleation of new particles via addition of more molecules to the existing clusters. This step is limited by diffusion, depending on the rate at which molecules can diffuse to the surface of the nucleus and attach themselves to it. Therefore, a key assumption of this model is that all particles that were nucleated at the same time grow at equal rates, resulting in the low polydispersity we observed in our PEG-DA formulations.

The ability to load and release IGF-1 from the 2 – 3 µm MPs developed here was tested. We found a direct correlation between AA content and loading efficiency, suggesting the presence of electrostatic interactions between the positively charged IGF-1 molecules and the electronegative carboxylate groups in the MPs. This trend was not found in the case for insulin loading, likely due to its positive charge at physiological pH not driving the same electrostatic binding. In the case of drug release, we observed a similar behavior with an inverse correlation between the release rate and the amount of AA in the MPs, with the total amount of IGF-1 released after 28 days decreasing from 94% to 71% when the PEG-DA was copolymerized with 80 mol% AA. Therefore, the release rate might be tuned by varying simple parameters in formulation, namely AA content and initial protein concentration. The utilization of varying levels of AA provides a degree of customization, allowing to tailor the microparticles’ charge based on the specific needs of the therapeutic molecule of interest. We also encapsulated IGF-1 loaded MPs within HA hydrogels, effectively adding another limiting factor for diffusion and further slowing down the release rate. The released IGF-1 retained its bioactivity after release up to 14 days as evidenced by DRG assays. Furthermore, the MPs exhibited very low cytotoxicity, presenting a promising platform to deliver growth factors.

While NT-3 and IGF-1 have both been found to be effective individually, the combined action of NT-3 and IGF-1 had not been extensively reported. After a literature search, very few instances of IGF-1 bioactivity assays with neurons were found ^44,45^. Here we found that the combination of optimal dosages of both growth factors had a synergistic effect on neurite outgrowth in mESC-derived V2a interneuron aggregates. The optimal concentration of NT-3 *in vitro* was previously determined, and in this work, we conducted a similar experiment with IGF-1. Individually, 20 ng/mL NT-3 and 50 ng/mL IGF-1 resulted in the highest neurite outgrowth, and the combination of both significantly increased the average neurite length. However, using half of the concentration for each growth factor (10 ng/mL NT-3 and 25 ng/mL IGF-1) did not decrease the growth, resulting in average lengths comparable to the higher concentration condition. This suggests that we can use smaller amounts of protein without affecting the overall axon-growth promoting effect. The retention of IGF-1’s bioactivity over extended periods post-loading, coupled with the observed synergistic effects with NT-3, underscores the potential of PEG-DA microparticles in enhancing the therapeutic impact of growth factors. The observed synergy could pave the way for dose optimization strategies, reducing potential side effects while maximizing therapeutic efficacy.

## Conclusion

In this work, we demonstrated the feasibility of using PEG-DA based microparticles as a drug delivery platform for growth factors and other proteins. We synthesized the MPs via precipitation polymerization in three different formulations, with varying levels of AA to alter the overall charge of the particles without significantly affecting the individual size and morphology. We found that through affinity binding, the growth factor IGF-1 could be loaded and released in a sustained manner over a period of 4 weeks. The release could be further slowed down by incorporating the particles within the HAmF scaffolds previously developed. The MPs showed very little cytotoxicity and the released IGF-1 retained its bioactivity up to two weeks post-loading, as shown by axonal growth assays in both primary DRG neurons and mESC-derived V2a interneurons. Furthermore, we found a synergistic effect between NT-3 and IGF-1 on neurite outgrowth, using lower than the optimal concentrations for each growth factor individually. The system reported here offers a versatile and scalable approach for the controlled delivery of bioactive molecules.

## Acknowledgements

This research was funded and supported by the National Institutes of Health (grant ID: R01-NS090617).

